# Swine influenza A virus isolates containing the pandemic H1N1 origin matrix gene elicit greater disease in the murine model

**DOI:** 10.1101/2023.09.25.559220

**Authors:** Shelly J. Curran, Emily F. Griffin, Lucas M. Ferreri, Constantinos S. Kyriakis, Elizabeth W. Howerth, Daniel R Perez, S. Mark Tompkins

## Abstract

Since the 1990’s, endemic North American swine influenza A viruses (swFLUAV) contained an internal gene segment constellation referred to as the triple reassortment internal gene (TRIG) cassette. In 2009, the H1N1 pandemic (pdmH1N1) virus spilled back into swine, but did not become endemic. However, the pdmH1N1 did contribute the matrix gene segment (pdmM) to the swFLUAVs circulating in the pig population; which replaced the classical swine matrix gene (swM) found in the TRIG cassette, suggesting that the pdmM has a fitness benefit. Others have shown that swFLUAV containing the pdmM have greater neuraminidase activity and transmission efficiency compared to viruses containing the swM gene segment. We hypothesized that the matrix gene could also affect disease and utilized two infection models, resistant BALB/c and susceptible DBA/2 mice, to assess swFLUAV pathogenicity. We infected BALB/c and DBA/2 mice with a panel of H1 and H3 swFLUAVs containing the swM or pdmM gene and measured lung virus titers, morbidity, mortality, and lung histopathology. H1 influenza strains containing the pdmM gene caused greater morbidity and mortality in both resistant and susceptible murine strains, while H3 swFLUAVs caused no clinical disease. However, both H1 and H3 swFLUAVs containing the pdmM replicated to higher viral titers in the lungs and pdmM containing H1 viruses induced greater histological changes compared to swM H1 viruses. While the surface glycoproteins contribute to swFLUAV pathogenicity, and other genes also influence disease, these data suggest that the origin of the matrix gene also contributes to pathogenicity of swFLUAV viruses in mice.

## INTRODUCTION

Influenza is considered a major public health threat causing between 250 and 300 thousand deaths annually worldwide (1-3). Influenza A viruses (FLUAVs) are single stranded, negative sense RNA viruses with a segmented genome, containing eight gene segments (4). These include the hemagglutinin (HA) and neuraminidase (NA) genes, which encode the surface glycoproteins forming the spikes on the outside of the virion and determine the virus subtype, the internal protein genes which include three polymerase, polymerase basic 1 (PB1), polymerase basic 2 (PB2), and polymerase acidic (PA) genes, as well as the nucleoprotein (NP), nonstructural (NS) and the matrix (M) segments (4).

The M gene encodes two proteins, M1 and M2. M2, is a 97 amino acid (aa), well-characterized proton channel, which contributes to the release of viral RNA from the virion during infection (5) as well as virus budding (6). The M2 proton channel also interacts with the host immune response modifying inflammasome activation (7) and autophagy (8-10). The 252 aa M1 protein bridges the viral ribonucleoprotein complexes (RNPs) and membrane proteins. It plays an important role in the release of viral RNA from the virion during infection, export of the viral RNPs out of the nucleus, inhibition of re-import of RNPs, and assembly of virus particles. The M1 protein is a determinant of the characteristic pleomorphic morphology of influenza virus particles, with filamentous virus particles linked to increased neuraminidase activity, affecting transmission (11-13).

Similar to humans, FLUAVs have been enzootic in swine worldwide, causing respiratory disease outbreaks in herds characterized by high morbidity, low mortality, and significant economic losses (14). For almost eight decades, a single H1N1 FLUAV, known as “classical” swine influenza, was circulating in the North American swine population (15, 16). However, in the late 1990’s novel triple reassortant H1N1, H1N2 and H3N2 viruses emerged in pigs containing gene segments from classical swine, human, and avian origin FLUAVs (17, 18). The six internal gene segments, known as the triple reassortment gene cassette (TRIG), became predominant; such that, since 2000 most fully characterized North American swine influenza strains contain this internal protein gene combination (16, 17, 19-22). Since the pandemic of 2009, which was caused by a novel reassortant swine-origin H1N1 virus with gene segments from North American swine viruses and an avian-like H1N1 Eurasian swine virus (23), the necessity for surveillance of the spread and genetic diversity of swine influenza viruses has become evident. Recent studies have shown regional differences in circulating strains both in the United States, as well as in other countries, with a diversification of genetic constellations and multiple HA and NA subtypes and clades in co-circulation (24-32). At the same time, field surveillance has demonstrated that since 2011, the pandemic origin M gene segment (pdmM) has been systematically replacing the swine TRIG M gene segment (swM) in North American swine FLUAVs, with up to 70% of swFLUAVs containing the pdmM gene in 2011 and increasing to 100% by 2015 (30, 33-35). The pdmM gene has been linked to increased neuraminidase activity and increased transmissibility (13, 36) and the pdmM gene was commonly found in swFLUAVs isolated at state fairs (37). Other studies have demonstrated that the M gene segment is under selective pressure independent of other genes with sites potentially related to host tropism and immune response (38, 39).

Predisposition to increased severity of disease and complications from FLUAV infection can be attributed to a variety of host factors including obesity, hypertension, and asthma, as well as host genetics (1, 40-42). The murine model is one of the most common animal models used for FLUAV pathogenesis research. Previous studies have shown a variation in pathogenesis between inbred strains of mice resulting in categorization of a continuum from susceptible to resistant (43). Susceptible strains, such as DBA/2, demonstrate higher morbidity, mortality, and viral replication while their immune response is characterized by greater concentrations of proinflammatory cytokines and increased numbers of lung infiltrates in response to influenza infection compared to resistant mouse strains such as BALB/c (43).

We hypothesized that infection of mice with swine FLUAV strains containing the pdmM gene would result in greater morbidity and mortality and induce more severe lung lesions compared to infection with strains of the same subtype and combination of gene segments containing swM gene. We used both DBA/2 and BALB/c mice to define the infection and disease phenotypes of swine influenza isolates to differentiate the contribution of the M gene in susceptible and resistant murine strains. We define virus replication kinetics, clinical disease, and pathology elicited by swM and pdmM gene containing viruses, demonstrating increased disease severity elicited by viruses with the pdmM gene.

## RESULTS

### The matrix gene contributes to morbidity and mortality of swine H1 influenza infection in mice

To reflect the potential for different morbidity and mortality outcomes dependent on the host, we inoculated both resistant (BALB/c) and susceptible (DBA/2) mouse strains (44) with a panel of swine H1 FLUAVs spanning a variety of HA and NA clades and containing either the swM or pdmM gene segment (n = 5 mice/group) (**Figure 1A**). The H1 viruses containing the pdmM caused greater morbidity, as demonstrated by weight loss, and mortality in both resistant (**Figure 1B and C**) and susceptible (**Figures 1D and E**) mouse strains. As previously shown by others, infection of mice with H3 viruses of human or swine origin resulted in minimal morbidity or mortality without adaptation (45, 46). To investigate whether the pandemic origin matrix gene can induce greater morbidity or mortality in mice infected with swine H3 influenza strains, we inoculated both resistant and susceptible murine strains with H3 influenza strains differing only in the matrix gene (n = 5 mice/group). Both resistant and susceptible mouse strains showed no morbidity, as determined by weight loss (**Figure 2**), or mortality (data not shown) when infected with H3 strains containing either the pdmM or swM gene segments.

**Figure 1.**
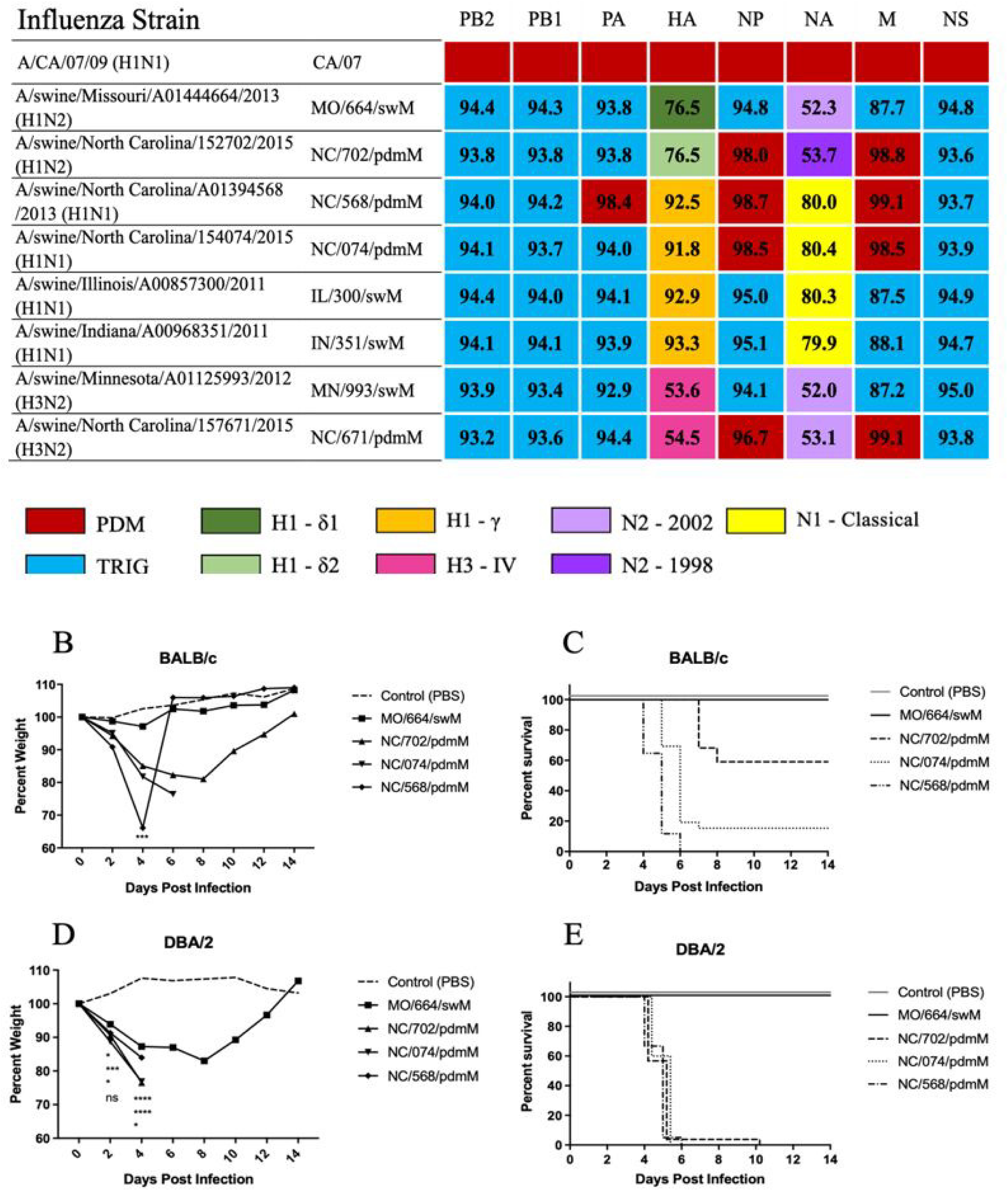
Influenza viruses used for mouse infections and morbidity and mortality in mice. (A) The origin of each gene segment is color coded according to the key. Numbers indicate % nucleotide sequence homology compared to A/CA/04/2009 (pdmH1N1). (B, D) Percent weight loss and (C, E) mortality of (B, C) BALB/c or (D, E) DBA/2 mice following infection with indicated swine H1 influenza viruses. Mice were inoculated with 1e5 pfu of virus (n=5/group) and weights were recorded every other day. This study was repeated to confirm results. Statistical comparison between MO/664/swM and the other viruses by two-way ANOVA with Dunnett *post-hoc* (B, D). * <0.05, **<0.005, *** <0.001, **** <0.0001. Kaplan-Meier survival curves (C, E). Abbreviations: PB2 polymerase basic 2; PB1 polymerase basic 1; PA polymerase acidic; HA hemagglutinin; NP nuclear protein; NA neuraminidase; M matrix; NS nonstructural; PDM pandemic 2009 lineage; TRIG triple reassortment internal gene constellation.

**Figure 2.**
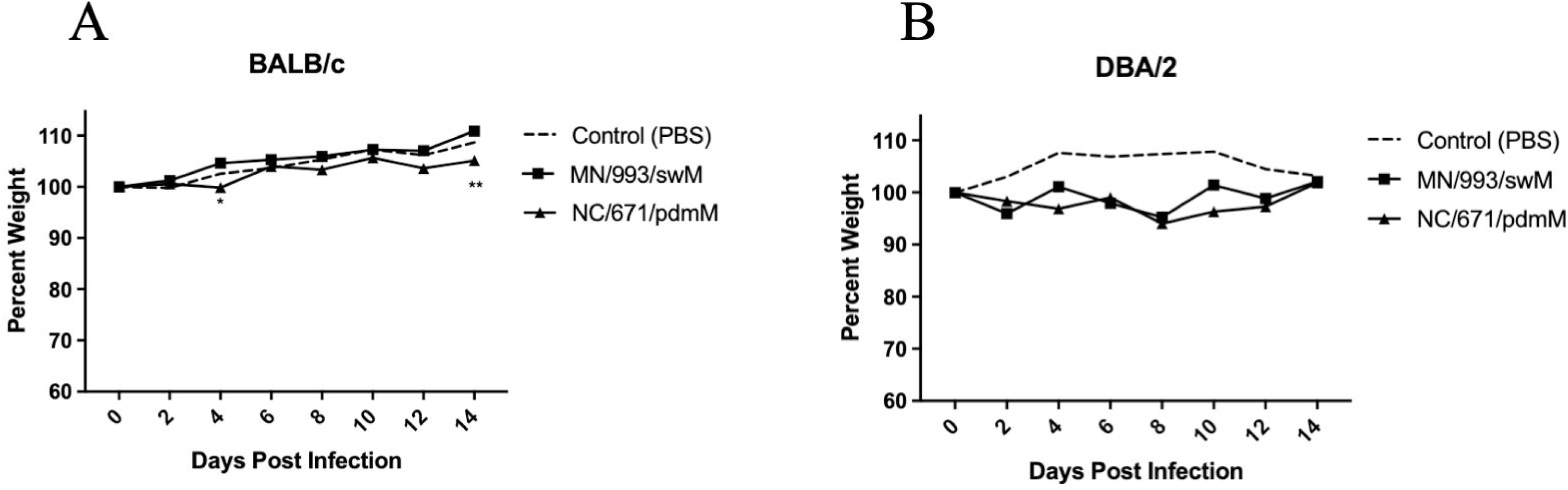
Morbidity of mice infected with swine H3 influenza viruses. (A) BALB/c or (B) DBA/2 mice were inoculated with 1e5 pfu of the indicated viruses (n=5 mice/group, repeated) and weights were recorded every other day. Statistical comparison between MN/993/swM and NC/671/pdmM by two-way ANOVA with Bonferroni *post-hoc*. **<0.005

### The matrix gene contributes to increased viral replication in the lungs of infected mice

Influenza-mediated morbidity and mortality in mouse models can be attributed to viral replication and immune mediated damage. In order to determine whether viral replication contributes to the greater morbidity and mortality caused by FLUAV strains containing the pdmM gene segment, we inoculated resistant and susceptible murine strains with swFLUAVs and assessed virus replication kinetics by lung virus titer (n = 5 mice/virus/collection timepoint). The H1 influenza viruses containing the pdmM gene segment replicated to significantly higher (between 1 and 4 logs greater) viral titers at 2 days post infection (DPI) and maintained significantly higher viral titers through 6 DPI in the resistant BALB/c mice (**Figure 3A**). The same viruses replicated only slightly better, 2 – 3 logs greater lung viral titers, than the swM- containing virus in the susceptible DBA/2 mouse strain, apart from NC/702/pdmM, which replicated similarly to the MO/664/swM virus (**Figure 3B**). Lung virus titers were also compared between H3 viruses containing pdmM or swM gene segments, in resistant and susceptible mouse strains. While neither of the swine H3 influenza viruses caused clinical signs, the viruses did replicate in mice, albeit at lower peak viral titers compared to the H1 swFLUAVs. Notably, NC/671/pdmM replicated to higher viral titers compared to MN/993/swM and was detectable through 6 DPI, while MN/993/swM was cleared after 4 DPI (**Figure 3C**). In the susceptible DBA/2 mouse strain, no significant differences were observed in virus replication kinetics, although titers following pdmM-containing H3N2 viral infection were consistently slightly higher until cleared after 6 DPI (**Figure 3D**). The robust replication of non-mouse-adapted H3N2 influenza viruses in mice was unexpected, so we assessed several human-origin H3N2 (huH3) viruses for replication in BALB/c and DBA/2 mice. While the swine H3N2 viruses replicated in mice without causing disease, huH3 viruses having HA and NA genes segments related to the swFLUAVs failed to replicate in either mouse strain, with no detectable virus in the lung at 2 DPI (data not shown). The greater lung viral titers of both H1 and H3 influenza strains containing the pandemic origin matrix gene has the potential to induce greater pathology in the lungs; therefore, we next assessed the lungs of influenza-infected mice for histopathologic changes.

**Figure 3.**
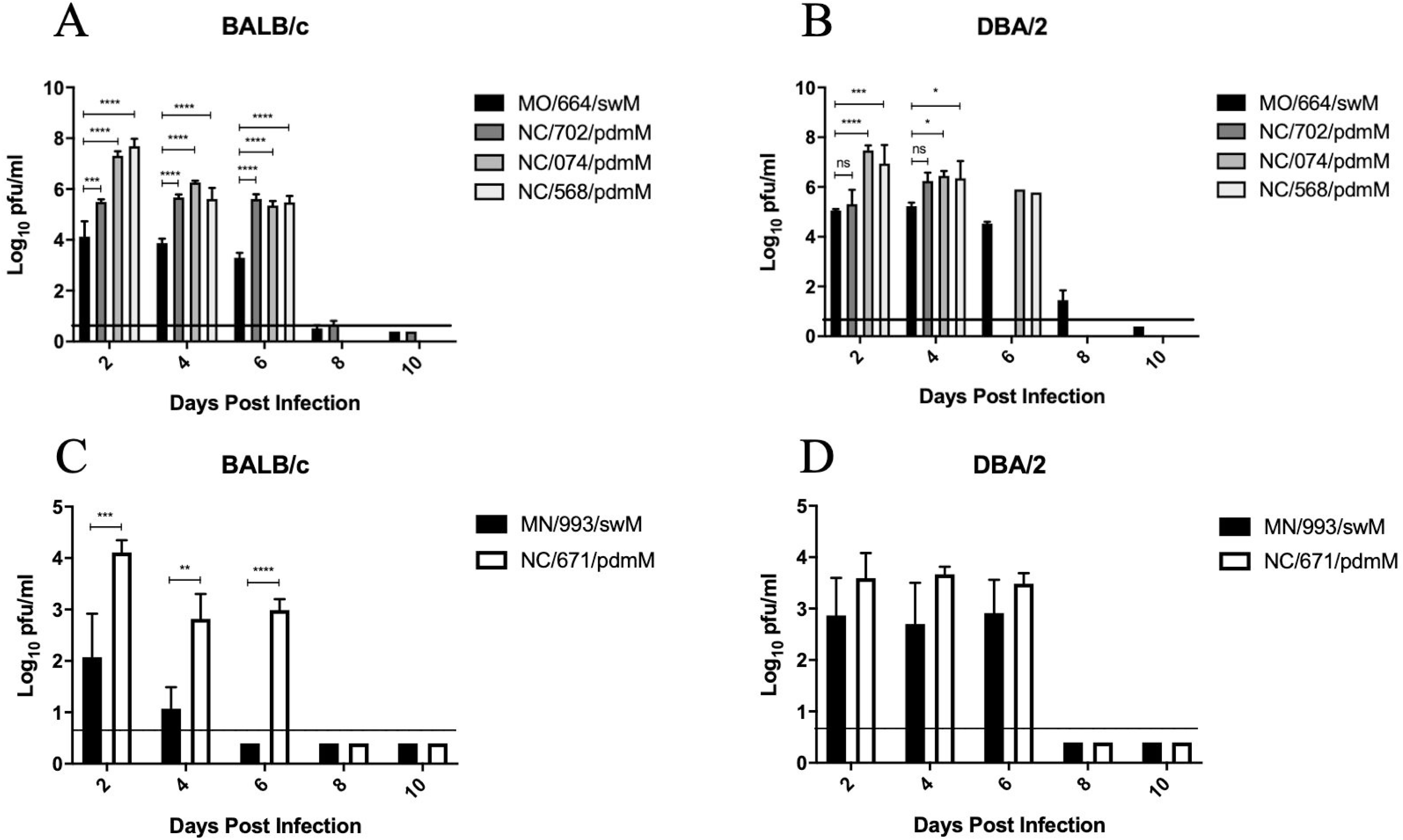
Lung virus titers over time from mice infected with swine H1 and H3 influenza viruses. (A, C) BALB/c and (B, D) DBA/2 mice were inoculated with 1e5 pfu of the indicated viruses. Lungs collected at indicated time points, homogenized, and assayed for virus titer by plaque assay (n=5 mice/group/collection timepoint, repeated). Statistical comparison between (A, B) MO/664/swM and the other viruses was completed using two way ANOVA with Dunnett *post-hoc* test or (C, D) MN/993/swM and NC/671/pdmM by two-way ANOVA with Bonferroni *post-hoc* test. * <0.05, **<0.005, *** <0.001, **** <0.0001.

### The pandemic origin matrix gene contributes to greater disease and more severe histological changes in the lungs of influenza-infected mice

In order to understand the development of disease and evaluate the characteristics and extent of lesions induced in the lung, we inoculated both resistant (BALB/c) and susceptible (DBA/2) mice with the panel of swFLUAVs and assessed histopathologic changes in the lung at 2 and 4 DPI (n = 2 mice/virus/collection timepoint). Mice inoculated with MO/664/swM had mild pulmonary changes on 2 DPI characterized by a small number of bronchioles with mild segmental necrosis of the epithelium and minimal peribronchiolar infiltrations of lymphocytes admixed with neutrophils. In addition, a small number of vessels with mild perivascular infiltrations of lymphocytes admixed with fewer neutrophils, and rare foci of alveoli containing small numbers of neutrophils and macrophages were present (data not shown). Changes were slightly more severe on 4 DPI (**Figure 4A, E**). The number of involved bronchioles and vessels increased, but was still less than 25%, and there was an increased amount of epithelial necrosis and numbers of peribronchiolar and perivascular inflammatory cells. Susceptible, DBA/2 mice were similarly infected with MO/664/swM and lungs analyses for pathological changes. Mild to moderate interstitial changes were present in the DBA/2 mice, characterized by mild multifocal alveolar infiltrations of a small number of neutrophils and macrophages (data not shown).

**Figure 4.**
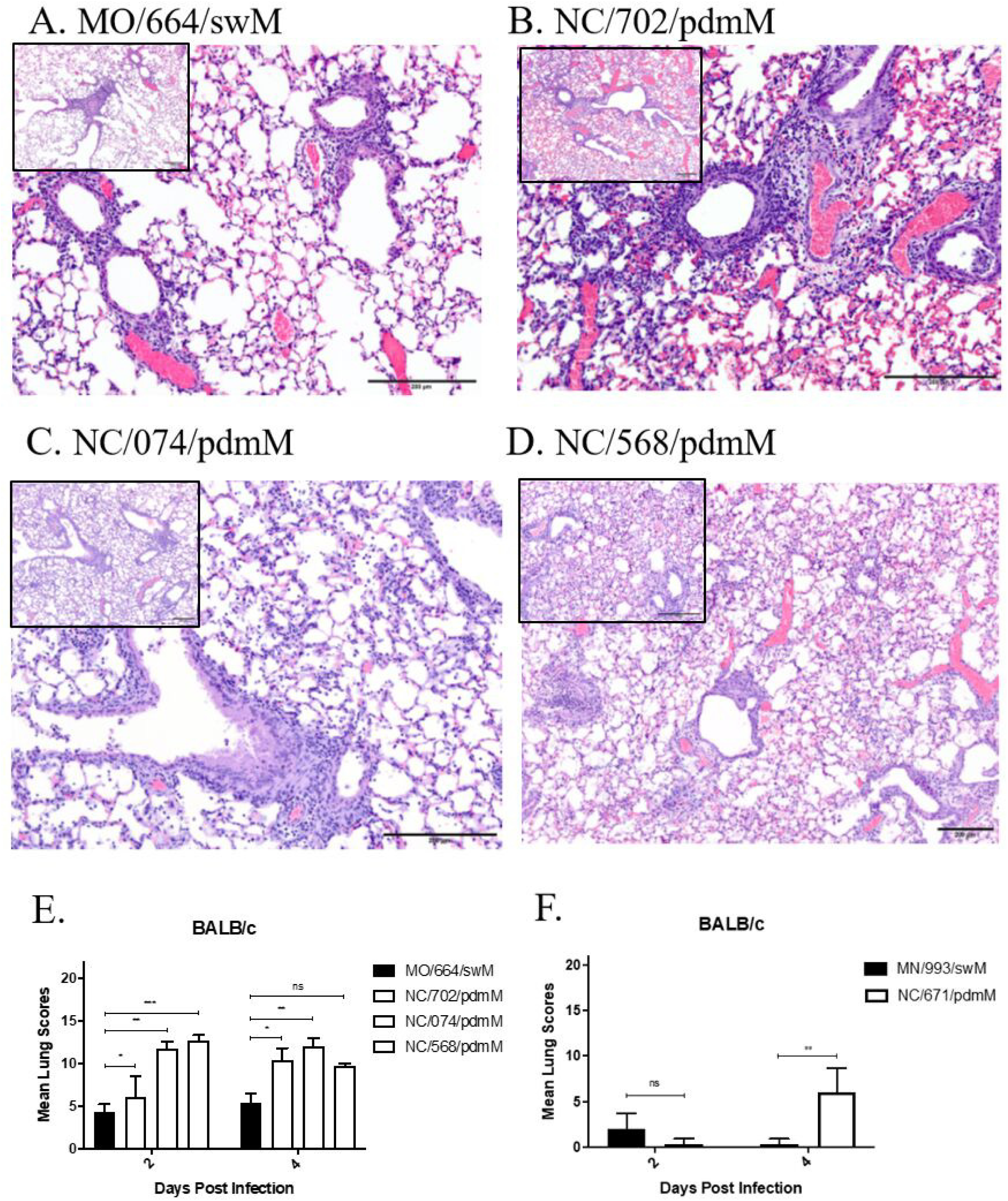
Histological images and lesion scores from mice infected with swine influenza viruses. BALB/c mice were inoculated with 1e5 pfu of the indicated viruses, euthanized, and lungs fixed for histological analysis at 2 and 4 DPI (n = 2 mice/virus/collection, repeated). Lungs were assigned histologic lesion scores out of 22. (A – D) Images of lung sections from 4 DPI at 20x and 10x (inset) for A/swine/Missouri/A01444664/2013 (H1N2) (A), A/swine/North Carolina/152702/2015 (H1N2) (B), A/swine/North Carolina/154074/2015 (H1N1) (C), and A/swine/North Carolina/A01394568/2013 (H1N1) (D). (A) MO/664/swM: Mild perivascular and peribronchiolar infiltrations of mostly lymphocytes and mild segmental necrosis of the bronchiolar epithelium are present, but there are no significant interstitial changes. Lesion score: 6/22. (B) NC/702/pdmM: Bronchioles are dilated and there are moderate peribronchiolar and mild perivascular infiltrations of mostly lymphocytes and diffuse necrosis of the bronchiolar epithelium. Focally extending from the central bronchiole, alveolar septa are thickened and there are small numbers of inflammatory cells in the alveoli. Lesion score: 10/22. (C) NC/074/pdmM: Bronchioles are slightly dilated with mild epithelial necrosis and sloughed epithelial cells and a few inflammatory cells in the lumen. There are mild to moderate peribronchiolar and perivascular infiltrations of mostly lymphocytes. Diffusely the alveolar septa are mildly thickened, and the alveoli contain small numbers of inflammatory cells. Lesion score: 13/22. (D) NC/568/pdmM: Bronchioles are slightly dilated and lined by attenuated epithelium. There are mild peribronchiolar and perivascular infiltrations of mostly lymphocytes. Diffusely the alveolar septa are mildly thickened, and alveoli contain small numbers of inflammatory cells admixed with erythrocytes. Lesion score: 10/22. (E and F) Histologic lesion scores out of 22 for BALB/c mice inoculated with either the H1 or H3 swine influenza isolates. Statistical differences were calculated using two-way ANOVA between (E) MO/664/swM and the other viruses with Dunnett *post-hoc* test or (F) MN/993/swM and NC/671/pdmM with Bonferroni *post-hoc* test. *<0.05, **<0.005, *** <0.001, **** <0.0001.

The resistant BALB/c mice inoculated with NC/702/pdmM, NC/074/pdmM, or NC/568/pdmM had similar pulmonary changes on 2 DPI compared to mice inoculated with MO/664/swM; however, histopathology was more severe, ranging from mild to moderate overall (**Figure 4E**), with larger numbers of involved bronchioles and vessels, more extensive epithelial necrosis, and increased numbers of perivascular and peribronchiolar lymphocytes admixed with neutrophils (data not shown). Pulmonary changes increased from mild to moderate in the NC/702/pdmM mice by 4 DPI (**Figure 4B**). However, in the NC/074/pdmM and NC/568/pdmM mice, changes were already moderate on 2 DPI, with the NC/568/pdmM inoculated mice being the most severe of all the groups. The resistant BALB/c mice inoculated with the pdmM-containing viruses had more extensive and severe interstitial involvement than seen in the swM-inoculated mice, which increased in extent and severity of involvement by 4 DPI, being most severe in mice inoculated with NC/568/pdmM (**Figure 4E**). In NC/702/pdmM and NC/074/pdmM inoculated mice, interstitial involvement ranged from mild (< 25%) on 2 DPI to moderate (25-50%) by 4 DPI and was characterized by foci with slightly thickened alveolar septa and small numbers of neutrophils, lymphocytes and macrophages in alveoli to foci where alveoli were filled with large numbers of inflammatory cells and necrotic debris (**Figure 4B and C**). In NC/568/pdmM inoculated mice, interstitial involvement was already moderate on 2 DPI (data not shown) with > 50% involvement of the parenchyma by 4 DPI (**Figure 4D**). Interstitial foci were similar to mice inoculated with NC/702/pdmM and NC/074/pdmM but also included foci with moderately thickened alveolar septa with mild epithelial hyperplasia. Mice inoculated with NC/568/pdmM by 4 DPI had multifocal interstitial hemorrhage that was more severe than with any of the other virus isolates (**Figure 4D**) and included the development of hyaline membranes in DBA/2 mice, which suggests more extensive alveolar septal damage with the NC/568/2013 isolate. Overall, there was significantly greater pulmonary changes in mice inoculated with swFLUAVs containing the pdmM gene segment compared to the virus containing the swM gene segment (**Figure 4E**). Similar results were observed in swFLUAV-infected, susceptible DBA/2 mice (data not shown).

Resistant and susceptible mouse strains were also inoculated with H3 swFLUAVs containing either the pdmM or swM gene and histopathologic changes were assessed in the lung at 2 and 4 DPI (n = 2 mice/virus/collection timepoint). On 2 DPI, lungs from BALB/c mice inoculated with MN/993/swM or NC/671/pdmM had minimal changes that were not specific for influenza, however by 4 DPI, mice inoculated with NC/671/pdmM had mild pulmonary changes consistent with influenza infection in contrast to minimal non-specific changes in MN/993/swM infected mice (**Figure 4F**). Pathological changes were overall similar in DBA/2 and BALB/c mice; however, pathology was slightly more severe in BALB/c mice and included minimal to mild lymphocytic infiltrations around a small number of bronchioles and vessels and minimal to mild segmental epithelial necrosis in a few larger apical bronchioles (data not shown). Together these data suggest that the pandemic origin matrix gene induces greater pathologic changes in the lung resulting in greater severity of disease in both susceptible and resistant murine strains. However, differences in H1 and H3 swFLUAVs suggest the HA and NA, as well as the overall gene composition can influence the pathogenicity of a particular influenza strain. To further address this possibility, we infected both resistant and susceptible strains with a panel of H1 influenza strains containing the swM gene but varying either in their HA or NA segments.

### The HA and NA gene segments can impact disease severity of swM gene containing viruses

To determine whether swine H1 influenza strains containing the swM gene with either HA or NA gene segments of different clusters can cause greater morbidity, mortality, and severity of disease, we inoculated both resistant and susceptible mouse strains with an alternate panel of FLUAVs with HA and NA gene compositions differing from the MO/664/swM strain (n = 5 mice/virus/collection timepoint). The swFLUAVs IN/351/swM and IL/300/swM contain HA gene segments from the gamma cluster and NA gene segments from the classical N1 (**Figure 1A**). Nucleotide and predicted amino acid sequence comparisons were analyzed to ensure high homology among PB2, PB1, PA, NP, and NS genes (≥ 91% nucleotide and ≥ 92% predicted amino acid homology) as well as within HA and NA clusters (**Figure 1A** and **S**. **Table 2**). In the resistant BALB/c mouse strain, the alternate H1 FLUAV strains containing the swM gene induced greater morbidity, as determined by weight loss, but not greater mortality (**Figure 5A**), despite replicating to significantly higher titers in the lung (**Figure 5B**). In the susceptible DBA/2 strain, the alternate H1 swM containing strains did cause both greater morbidity and mortality without replicating to significantly higher titers (**Figure 5F – H)** unlike the pdmM containing viruses seen previously. Due to the potential for the increased replication of IN/351/swM and IL/300/swM to induce greater pathology, we evaluated the lungs for histopathologic changes (n = 2 mice/virus/collection timepoint).

**Figure 5.**
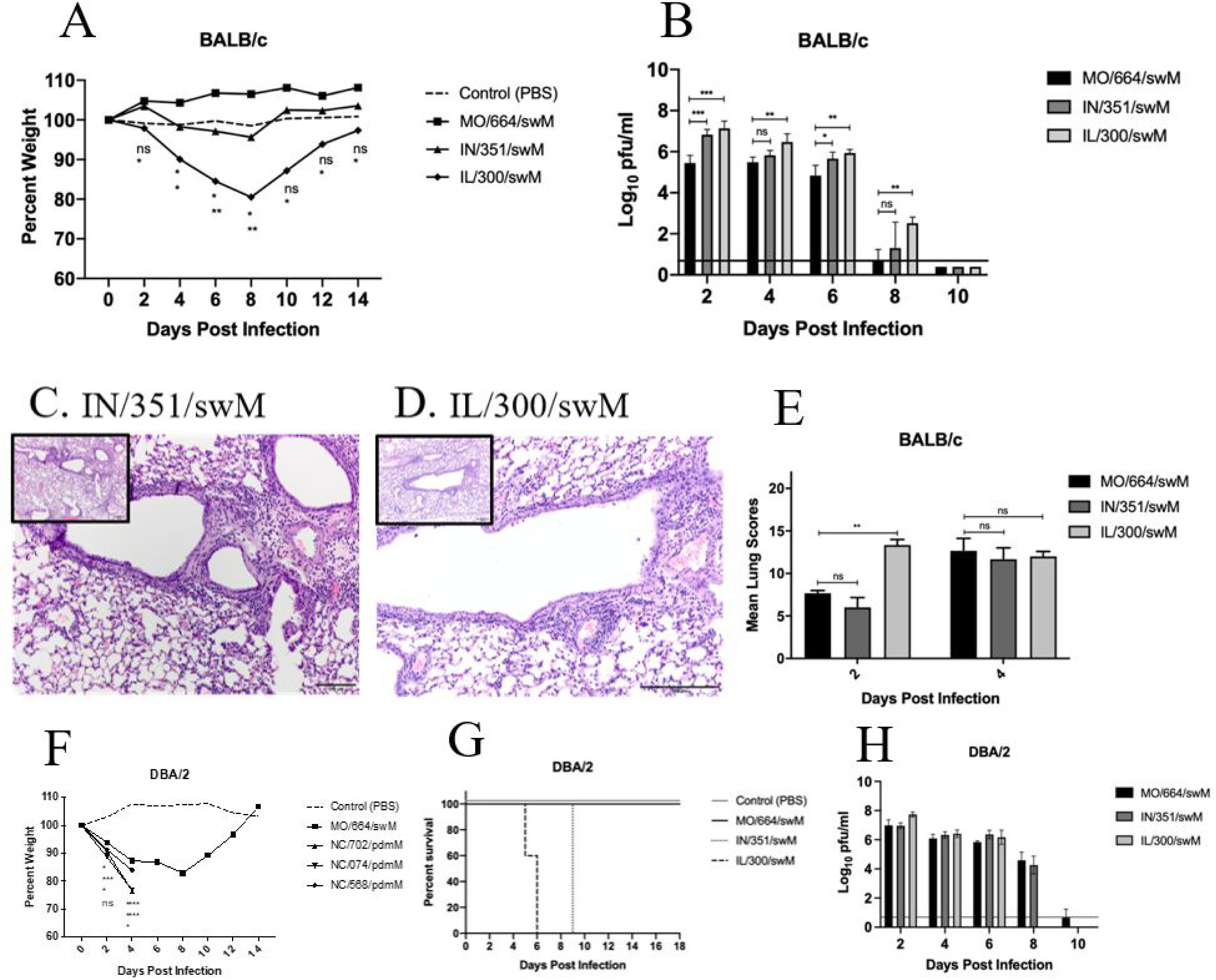
Morbidity, mortality, histology, and lung virus titers of mice infected with alternate swine H1 influenza viruses. (A – E) BALB/c mice were inoculated with 1e5 pfu of the indicated viruses and (A) weights were recorded every other day and lungs collected and (B) assayed for virus titer or (C, D, E) histopathology (n = 5 mice/virus/collection for titers, n = 2 mice/virus/collection for histopathology, repeated). Images of lung sections from (C) A/sw/Indiana/A00968351/2011 (H1N1, IN/351/swM) or (D) A/sw/Illinois/A00857300/2011 (H1N1, IL/300/swM) infected mice, 4 DPI at 20x and 10x (inset) and (E) histologic lesion scores out of 22. DBA/2 mice were inoculated with 1e5 pfu of the indicated viruses and (F) weights and (G) survival recorded, or (H) lungs collected and assayed for virus titer. Statistical differences were calculated between MO/664/swM and the other viruses by two-way ANOVA with Dunnett *post-hoc* test. * <0.05, **<0.005, *** <0.001, **** <0.0001.

For both murine models, on day 2, the predominant lesion was in the bronchioles characterized by epithelial necrosis and minimal interstitial changes consistent among the three swFLUAVS MO/664/swM, IN/351/swM, and IL/300/swM (data not shown). By day 4, in both the resistant BALB/c (**Figure 5C and D**) and susceptible DBA/2 strains (data not shown), there were no significant differences in lung pathology scores for the three swM gene containing viruses (**Figure 5E**). Vasculitis was present with all three swFLUAVs in BALB/c and DBA/2 mice (data not shown). Overall, there were minimal pulmonary differences in the lungs of mice infected with any of the swFLUAVs containing the swM gene, in contrast to the significant pulmonary changes observed between mice infected with the swM versus the pdmM gene.

## DISCUSSION

Multiple introductions of the 2009 pH1N1 virus into the swine population has led to reassortment events with previously circulating swine FLUAVs resulting in hitherto unseen genetic constellations. Prior research has shown the matrix gene from the pandemic virus is replacing the classical swine origin matrix gene found in the TRIG cassette, suggesting an evolutionary advantage or fitness benefit. We used resistant and susceptible mouse strains to investigate whether infection with swine influenza isolates containing the pandemic origin matrix gene (pdmM) would induce greater disease compared to isolates containing the classical swine origin matrix (swM) gene. We chose influenza isolates reflecting the predominant strains found in North America between 2010 and 2016 (33). Infection with H1 influenza viruses containing the pdmM resulted in greater morbidity and mortality in both susceptible and resistant mouse strains. However, infection with H3 isolates containing either the pdmM or swM did not induce morbidity or mortality, which is in agreement with previous studies that have shown a very high lethal dose for earlier swine H3 viruses and an inability to infect mice with human H3N2 strains (46).

We evaluated virus replication in the lungs of both susceptible (DBA/2) and resistant (BALB/c) mice. Both H1 and H3 swFLUAVs containing the pdmM replicated to higher viral titers in the lungs of susceptible and resistant mice compared to swFLUAVs containing the swM. Higher viral titers in the mouse lungs suggests that the pdmM gene confers increased infectivity or replication efficiency within the mice at least to some degree. The higher viral load in the lungs may contribute to the greater morbidity and mortality observed in pdmM-containing influenza virus-infected mice. However, in comparing several swM-containing viruses, we observed varying lung virus titers in BALB/c mice without concomitant differences in mortality, and in contrast, no differences in lung virus titer yet clear differences in survival in susceptible DBA/2 mice. Thus, while the M gene may affect outcomes of infection, other gene segments also impact infection and disease, and is additionally influenced by the host genetic background. These differences could be due to virus replication at the cellular level or the host immune response to infection.

Others have compared outcomes of influenza virus infection in susceptible versus resistant mouse strains with avian H5N1 (47), mouse-adapted H1N1 (A/PR/8/1934) (48) or H3N2 (A/HKX-31) (49), or mouse-adapted classical swine H1N1 (50) virus. Influenza viruses replicated to higher titers in susceptible, DBA/2 mice compared to resistant (BALB/c or C57Bl/6) mice in all of these studies, and was correlated with increased inflammatory responses as well as increased disease severity and mortality. In contrast, virus titers were imperfectly correlated with morbidity and mortality in DBA/2 and BALB/c mice when infected with swFLUAVs containing swM or pdmM genes. The H1 influenza viruses containing the pdmM gene replicated to similarly high titers in BALB/c and DBA/2 mice, resulting in greater morbidity and mortality compared to the swM gene containing H1 virus, MO/664/swM, that replicated to lower virus titers and showed no mortality in either mouse strain. However, two other swM gene-containing H1 viruses replicated to higher titers in BALB/c mice, but did not cause mortality. Further the three swM H1 viruses, MO/664/swM, IN/351/swM, and IL/300/swM replicated to similar titers in DBA/2 mice with variable mortality. Finally, the tested H3N2 swFLUAVs replicated to similar or slightly lower titers compared to the H1N2 MO/664/swM virus, with the swM-gene containing H3N2 virus replicating more poorly than the pdmM virus, but neither caused morbidity or mortality in BALB/c or DBA/2 mice. While total virus replication or peak viral load may contribute to disease severity between susceptible and resistant mouse strains, other factors may be affecting disease severity.

Differences in the pro-inflammatory response between susceptible and resistant mouse strains has also been implicated in disease severity and mortality, with increases in pro-inflammatory cytokines and other inflammatory mediators observed in susceptible, DBA/2 mice associated with increased disease severity and mortality following influenza virus infection. We did not assess cytokine responses here; however, histopathological changes were generally greater in DBA/2 mice compared to BALB/c mice, suggesting increased inflammation.

The influenza matrix gene has been shown to influence the inflammasome and autophagy (7-10) as well as affect morphology, host range, and transmission (13, 38, 51). In humans, inflammation and development of pneumonia has been correlated with swine-origin, pandemic H1N1 infection (52), while other studies have related an altered immune response with FLUAVs containing the pdmM gene (53). We assessed the development of microscopic changes in the lungs of swFLUAV-infected BALB/c and DBA/2 mice. Both H1 and H3 swFLUAVs containing the pdmM gene induced greater histologic changes, characterized by necrosis, increased infiltrates, thickening of the alveoli septa, and epithelial hyperplasia compared to viruses containing the swM gene. This was seen in both resistant and susceptible mouse strains. The increased severity of lesions in the lungs of mice infected with an H3 swFLUAV containing the pdmM gene compared to the H3 virus containing the swM, while not as severe as was seen with the H1 isolates, was evident despite a lack of significant difference in viral replication in the susceptible murine strain. Together these data suggest that while greater viral replication by the swFLUAVs containing the pdmM gene contributes in part to the increased histopathologic lesions in the lungs, other viral and host factors may also contribute to the enhanced lung lesions and subsequent disease. The lack of significant pulmonary changes in the lungs of mice infected with the panel of swFLUAVs containing the swM gene despite differences in replication supports this concept. Of note, the microscopic changes in the lung switched from bronchiolar to more interstitial over time in the mice inoculated with the H1 viruses containing the pdmM gene. In humans, interstitial damage along with the development of hyaline membranes has been associated with the development of severe influenza viral pneumonia (54). Furthermore, in mice as well as in humans, severe influenza viral pneumonia has been correlated to the dysregulation of inflammation of the airways (55-58). This suggests that the differential lung pathology may in part be due to the subsequent immune response to the infection, and the potential for infection with swine influenza isolates containing the pandemic matrix gene to generate an exacerbated immune response in the murine model.

Studies using reverse genetics viruses would be useful in ascertaining to what degree the pandemic matrix gene influences virus replication, disease progression, and outcome in murine models. Others have shown the origin of viral segments such as PB1 and NS1 genes may also contribute to swine influenza strain virulence and disease (59, 60). Recently, a reverse genetics approach was used to compare influenza virus replication and disease in BALB/c mice, comparing a Eurasian-origin M gene with the pdmM gene in an Asian-origin H1N1 virus background (61). The pdmM gene conferred increased virus titers, mortality, and disease, supporting our hypothesis and observations, although the Eurasian M gene is distinct from the classical swM gene we assessed here (16). Thus, assessment of the role of the M gene in our model by reverse genetics will be important. Others have assessed immune responses in susceptible and resistant mouse strains infected with avian, human (mouse-adapted human), and swine origin FLUAVs, noting increased proinflammatory immune responses in susceptible mouse strains (47-50). Understanding differences in the immune response elicited by swM- compared to pdmM- containing swFLUAVs would further clarify mechanisms of increased disease in the different mouse strains and in the context of the matrix gene. These studies along with the work reported here will help define the impact of individual genes on virus replication, the host response to infection, and determinants of disease.

## MATERIALS AND METHODS

### Cell Culture and Virus Propagation

Influenza viruses A/swine/Missouri/A01444644/2013 (H1N2; MO/664/swM), A/swine/North Carolina/A01394568/2013 (H1N1; NC/568/pdmM), A/swine/Minnesota/A01125993/2012 (H3N2; MN/993/swM), A/swine/Illinois/A00857300/2011 (H1N1; IL/300/swM), and A/swine/Indiana/A00968351/2001 (H1N1; IN/351/swM) were obtained from the USDA National Veterinary Services Laboratories (NVSL) reagent resource (30). Influenza viruses A/swine/North Carolina/152702/2015 (H1N2; NC/702/pdmM), A/swine/North Carolina/154074/2015 (H1N1; NC/074/pdmM), and A/swine/North Carolina/157671/2015 (H3N2; NC/671/pdmM) were obtained through the swine influenza surveillance project of the NIAID Emory-UGA Center of Excellent in Influenza Research and Surveillance (CEIRS) (28, 62). Viruses were propagated in Madin-Darby canine kidney (MDCK) cells (IRR FR-926) in Minimal Essential Media [MEM (GIBCO™)] with 0.002μg tosyl phenylalanyl chloromethyl ketone (TPCK) trypsin (Worthington). Viruses were cultured to achieve stock titers above 1e6 pfu/ml. Influenza virus titers were assessed by plaque assay on MDCK cells as previously described (63). Briefly, a 24-well plate of MDCK cells was incubated with serial dilutions of virus at 37°C and 5% CO_2_ for 1-2 hours. The supernatant was removed, 1ml of 1:1 2.4% Avicel solution and overlay [MEM (GIBCO™) with 1M HEPES (GIBCO™), 200mM GlutaMAX-I, 7.5% NaHCO_3_ (GIBCO™), and antibiotic/antimycotic (GIBCO™)] was added and the cells were incubated at 37°C and 5% CO_2_ for 48 -72 hours prior to fixation with 80/20 methanol/acetone and staining with crystal violet.

### Sequencing and Analysis

For the sequencing of viral gene segments, viral RNA was isolated using RNAzol®RT (Sigma-Aldrich) as per manufacturer’s protocol. cDNA synthesis and PCR were performed using SuperScript III One-step RT-PCR (Invitrogen) as per manufacturer’s protocol. Primers used were as follows: All 8 genes simultaneously MBTUni-12 (Forward) 5’ACGCGTGATCAGCRAAAGCAGG3’, MBTUni-13 (Reverse) 5’ACGCGTGATCAGTAGAAACAAGG3’, PB2 Forward 5’AGCRAAAGCAGGTCAATTATATTCA3’, PB2 Reverse 5’AGTAGAAACAAGGTCGTTTTTAAACTA3’, PB1 Forward 5’AGCRAAAGCAGGCAAACCATTTGAATG3’ PB1 Reverse 5’AGTAGAAACAAGGCATTTTTTCATGAA3’, PA Forward 5’ AGCRAAAGCAGGTACTGATYCGAAATG3’, and PA Reverse 5’AGTAGAAACAAGGTACTTTTTTGGACA3’. NGS was performed using the Illumina MiSeq platform. All sequences are publicly available in GenBank. Virus isolates sequenced for this study are listed under NCBI bioproject accession number PRJNA600894 (**S. Table 1**). Sequences were aligned using MUSCLE alignment and comparison of predicted amino acids using Geneious (Biomatters Ltd.). Sequences of each of the genes of the isolates were compared to reference genes of pandemic origin, TRIG origin, and classical swine origin. Genes were categorized by the highest percentage similarity between the isolate gene and the reference sequence (**S. Table 2**).

### Illumina MiSeq Sequencing Platform

#### Amplicon purification and library preparation

The PCR products were purified using Agencourt AMPure XP Magnetic Beads (Beckman Coulter) at 0.45× and eluted in 30 µl of HyClone molecular biology water (Genesee Scientific). The concentrations of the eluates were measured using the Qubit dsDNA HS Assay kit (ThermoFisher) on the Qubit 3.0 fluorometer (ThermoFisher). Normalization was done at 0.2 ng/µl. Adapters were added using the Nextera XT DNA library preparation kit (Illumina) with 40% of the suggested final volume. The libraries were cleaned using 0.7× Agencourt AMPure XP Magnetic Beads, and the fragment size distribution was evaluated on the Agilent Bioanalyzer using the High Sensitivity DNA kit (Agilent). Thereafter, the samples were normalized to 4 nM. The pooled libraries were loaded at a concentration of 15 pM and sequenced using the MiSeq v2, 300 cycle reagent Kit (Illumina) in a paired-end fashion (150 × 2).

#### Genome assembly

Genome assembly was performed using a pipeline developed previously by Harm Van Bakel from Icahn School of Medicine at Mount Sinai (64). Initially, Cutadapt was used to remove low-quality sequences and adapters from paired fastq files. The initial assembly was conducted using the inchworm module of Trinity (65). Viral contigs containing internal deletions were detected by mapping against nonredundant IRD reference sequencing using BLAT (66). Afterwards, the inchworm assembly was iterated to eliminate breakpoint-spanning kmers. The resulting contigs were then oriented and trimmed, removing low-coverage ends as well as extraneous sequences beyond the conserved FLUAV termini. Finally, to evaluate assembly contigs and contiguity for all segments, sequences reads were mapped back to the ultimate assembly using Burrows-Wheeler Alignment (67).

### Mice

Female 6 – 8-week-old BALB/c mice were purchased from Charles River Laboratories (Raleigh, NC). Female, 6 – 8-week-old DBA/2 mice were purchased from Jackson Laboratories (Bar Harbor, ME). All animal studies were approved by the Animal Care and Use Committee of the University of Georgia and carried out in strict accordance with the National Institutes of Health Guide for the Care and Use of Laboratory Animals. Humane euthanasia of mice followed American Veterinary Medical Association guidelines. All murine experiments were repeated to confirm findings and increase robustness.

### *In vivo* infection

Mice were anesthetized by isoflurane (Patterson Veterinary) inhalation and intranasally inoculated with 50μl virus diluted in phosphate buffered saline (PBS). Control mice were inoculated with 50μl PBS.

### Lung virus titers

Mice were humanely euthanized, lungs were collected at set time points, homogenized in 1 ml cold PBS, centrifuged, and supernatant was aliquoted, and frozen at -80°C. Individual lung homogenate aliquots were thawed and assayed for influenza virus by plaque assay on MDCK cells.

### Histopathology

Mice were humanely euthanized at 2 and 4 DPI and lungs were collected, inflated and fixed by immersion in 10% buffered formalin. Fixed lungs were submitted to the American Association of Veterinary Laboratory Diagnosticians accredited histology laboratory in the Department of Pathology at the University of Georgia for processing and sectioning. Briefly, lungs were embedded in paraffin so that all lung lobes could be evaluated, and 4 μm sections were cut and stained with hematoxylin and eosin. Lungs were blinded, analyzed by a Diplomate of American College of Veterinary Pathologists, and scored as follows: Perivascular (bronchial tree) inflammation (0=none;1=mild, 1-2 cells wide; 2=moderate, 3-10 cells wide; 3=severe, >10 cells wide); percentage of bronchioles affected (0=none; 1=<25%; 2=25-75%; 3=>75%); peribronchiolar inflammation (0=none; 1=mild; 2=moderate; 3=severe); severity of airway luminal exudate, epithelial necrosis and inflammation (0=none; 1=mild; 2=moderate; 3=severe); percentage of alveolar involvement (0=none; 1=<25%; 2=25-50%; 3=>50%); severity of interstitial inflammation (0=none; 1=mild; 2=moderate; 3=severe); edema (0=none; 1=present); hemorrhage (0=none; 1=present); type II cell hyperplasia (0=none; 1=present);vasculitis (0=none; 1=present). This resulted in scores ranging from 0 - 22. BALT (well defined aggregates of mixed lymphocytes or follicles) was scored separately as 0-3 (0=none; 1=mild; 2=moderate; 3=severe).

### Statistical Analysis

Statistics were run using GraphPad Prism version 7.03. Statistical analysis included two-way analysis of variance (ANOVA) with Bonferroni *post-hoc* for weight loss, viral titers, and lung pathology scores of H3 swFLUAVs, Dunnett *post-hoc* for weight loss, viral titers, and lung pathology scores of H1 swFLUAVs or Kaplan-Meier Survival Curve for survival data. All results were considered significant at p-values < 0.05.

## ACKNOWLEDGEMENTS

We thank the staff of the UGA Animal Health Research Center and Animal Resources Program for their care and maintenance of the mice. We sincerely appreciate the Histology Laboratory in the Department of Pathology at the University of Georgia, specifically Dr. Elizabeth Howerth for lung histopathological analysis. Further, we are grateful to Cheryl Jones and Scott Johnson for their kind assistance in data collection. Madin-Darby Canine Kidney (MDCK-ATL) Cells, FR- 926, were obtained through the International Reagent Resource, Influenza Division, WHO Collaborating Center for Surveillance, Epidemiology and Control of Influenza, Centers for Disease Control and Prevention, Atlanta, GA, USA.

This work was supported by the UGA-Emory Centers of Excellence for Influenza Research and Surveillance contract HHSN272201400004C (S.M.T.).

Conceptualization, S.J.C., C.S.K., and S.M.T.; Methodology, S.J.C., L.M.F., E.F.G., and S.M.T., Formal Analysis, S.J.C., L.M.F., and E.F.G.; Investigation, S.J.C., and E.F.G.; Resources, S.M.T., E.W.H., and D.R.P.; Writing – Original Draft: S.J.C.; Writing – Review & Editing: S.J.C., E.F.G., S.M.T., L.M.F., C.S.K., E.W.H., and D.R.P.; Visualization, S.J.C., E.F.G., and S.M.T.; Supervision, D.R.P. and S.M.T.; Funding Acquisition, S.M.T.. All authors have approved the manuscript for publication.

We declare no conflict of interest.

## Data Availability

All sequences are publicly available in GenBank. Virus isolates sequenced for this study are listed under NCBI bioproject accession number PRJNA600894.

