## Supplemental Tables 1 and 2 for "Swine influenza A virus isolates containing the pandemic H1N1 origin matrix gene elicit greater disease in the murine model"

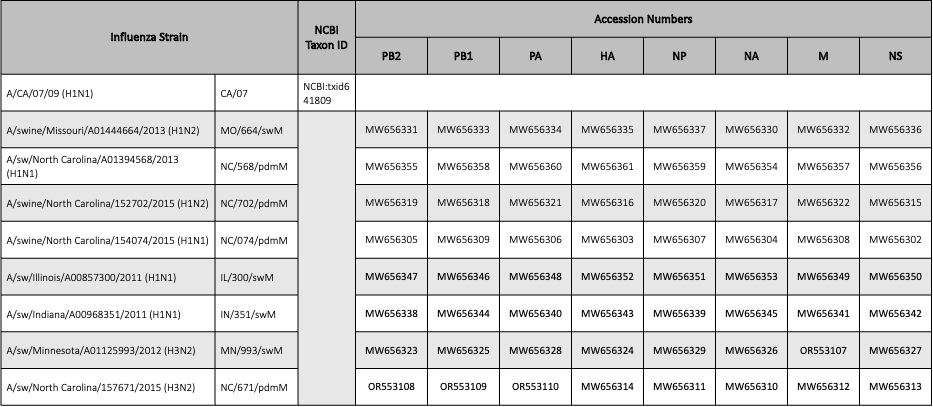

**S. Table 1 NCBI taxonomy ID and GenBank accession numbers for the virus strains used in this study.**

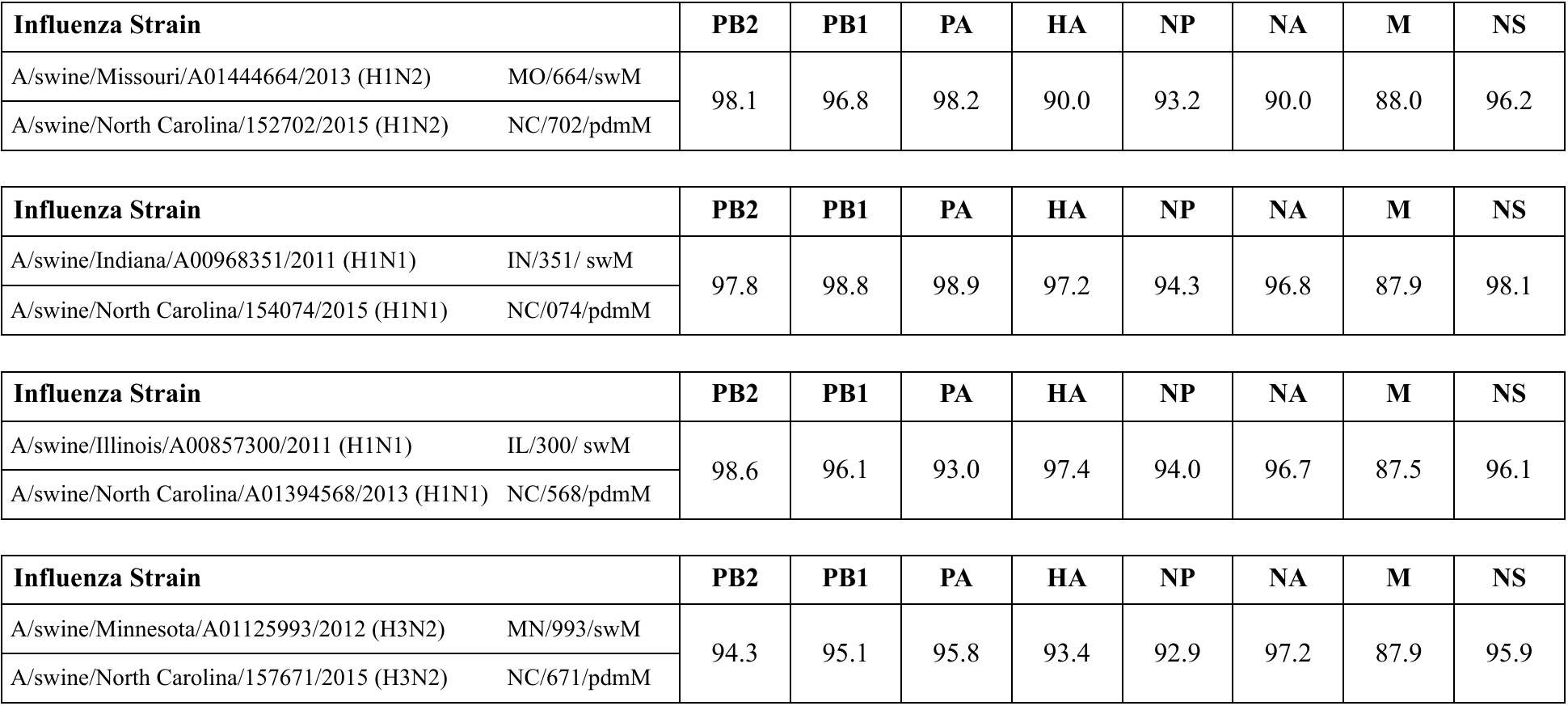

**S. Table 2. Percent homology based on predicted amino acid sequence comparison between matching gene segments**
